# Adoptive transfer of allogeneic gamma delta T cells promotes HIV replication in a humanized mouse model

**DOI:** 10.1101/2021.02.08.430263

**Authors:** Shivkumar Biradar, Yash Agarwal, Michael T. Lotze, Charles R. Rinaldo, Moses T. Bility, Robbie B. Mailliard

**Affiliations:** Department of Infectious Diseases and Microbiology, University of Pittsburgh, Graduate School of Public Health, Pittsburgh, PA, USA; Department of Surgery, University of Pittsburgh School of Medicine, Pittsburgh, PA, USA; Immunology, University of Pittsburgh School of Medicine, Pittsburgh, PA, USA; Bioengineering, University of Pittsburgh School of Medicine, Pittsburgh, PA, USA; Pathology, University of Pittsburgh School of Medicine, Pittsburgh, PA, USA; Microbiology and Molecular Genetics, University of Pittsburgh School of Medicine, Pittsburgh, PA, USA

## Abstract

Gamma-delta (γδ) T cells recognize antigens in an MHC-independent manner, with demonstrable cytotoxicity against cancer and virally infected cells. Human immunodeficiency virus (HIV) infection severely depletes the Vγ9Vδ2 (Vδ2) subset of these T cells in most infected individuals, with the exception of elite controllers. The capacity of Vδ2 cells to kill HIV-infected targets has been demonstrated *in vitro*, but this has not been verified *in vivo*. Here, we examined the immunotherapeutic potential of Vδ2 cells in controlling HIV replication *in vivo* and provide the first characterization of reconstituted γδ T cell subsets in the peripheral blood and lymphoid tissue in a humanized mouse model. We demonstrate the depletion of Vδ2 cells and increase in Vδ1 cells in the blood following HIV infection, similar to that observed in HIV-infected humans. The functionality of human Vδ2 cells isolated from humanized mice was confirmed via *ex vivo* expansion in response to zoledronate and IL-2 treatment. The adoptive transfer of activated Vδ2 cells failed to control HIV infection *in vivo* but instead exacerbated viremia by serving as early targets for HIV infection. Our findings suggest that Vδ2 cells play a critical and unappreciated role as early HIV targets of infection to promote viral dissemination.

## Introduction

Human gamma-delta (γδ) T cells mediate potent antiviral effects in an MHC-independent manner, recognize stressed cells similar to that of NK cells, and are widely distributed throughout barrier tissues [1-3]. Although human γδ T cells typically make up <10% of the total T cell population, they play an important role in recognizing nonpeptide antigens from various microbes, particularly malaria [4], and provide a mixture of innate and adaptive immune responses [5]. HIV infection has a dramatic and immediate impact on γδ T cells, inverting the normal proportions of the two major subsets of γδ T cells (designated Vδ1 and Vδ2) by selectively depleting Vδ2 T cells expressing the phosphoantigen-responsive Vγ9 chain (Vγ9Vδ2 T cells) [6]. Natural history studies of HIV infection demonstrate an inverse correlation between Vδ2 T cell frequency and HIV viral titers [7]. Earlier clinical reports demonstrated that, unlike most HIV-infected individuals, Vδ2 T cells are maintained at a normal frequency in elite controllers [7]. The hypothesis for these observations is that γδ T cells may provide protective immunity against HIV infection by secreting chemokines that compete for HIV entry coreceptors or by recruiting and promoting the activity of other immune cells to eliminate infected targets. A few *in vitro* studies demonstrated the cytotoxic capacity of Vδ2 T cells against HIV-infected targets [8, 9], but their *in vivo* function and therapeutic potential in HIV infection have yet to be fully elucidated.

Non-human primate models of simian immunodeficiency virus (SIV) dominate the current *in vivo* approaches to understanding the relationship between HIV viremia and γδ T cells. SIV contains only about 50 percent of the genetic code of HIV and there are substantial differences in γδ subset composition and phenotype in monkeys and humans [10]. The information we can extrapolate from non-human primate models of SIV becomes limited by the unaltered peripheral Vδ1/Vδ2 T cell ratio in SIV-infected macaques [11] and the genetic differences between SIV and HIV [10]. Therefore, an alternate approach is needed to understand the *in vivo* dynamics of γδ T cells in HIV infection. Among the widely used small mammal platforms for investigating HIV pathogenesis and therapeutics is the mouse model utilizing bone marrow-liver-thymus (BLT) humanized mice (huMice). Generated via peripheral injection of CD34+ hematopoietic stem cells (HSCs) and autologous transplantation of fetal liver and thymic explants into immunodeficient mice, BLT huMice provide both the peripheral immune circulation and human lymphoid microenvironment to study HIV in blood and human lymphoid tissues. Previously it has been shown that human CD4^+^/CD8^+^ T cell ratios before and after HIV infection of BLT huMice are comparable to clinical values seen in natural human infection [12]. While Vδ2 T cells become depleted during early stages of natural infection, often before the CD4^+^/CD8^+^ T cell ratio inverts, the impact of HIV infection on γδ T cells has yet to be fully characterized in the huMouse model.

In the present study, we provide the first reported phenotypic and functional characterization of human γδ T cells in BLT huMice and evaluate how they are impacted by HIV infection *in vivo*, and we assess their therapeutic potential following adoptive cell transfer. We demonstrate that the BLT huMouse model recapitulates the clinical changes in Vδ1 and Vδ2 T cell frequencies in the peripheral blood reported during natural HIV infection, providing for the first time an *in vivo* model relevant for studying human γδ T cell biology. When tested therapeutic impact on HIV infection, the adoptive transfer of cultured allogenic Vδ2 T cells into huMice implanted with CD4^+^ T cells from HIV-infected individuals on ART surprisingly resulted in enhancement rather than control of HIV replication. This escalation in viral production was accompanied with a marked increase in detection HIV p24 positive Vδ2 T cells isolated from the mice following adoptive transfer, suggesting that the Vδ2 T cells served as targets for HIV infection.

## Results

### Reconstitution of human γδ and αβ T cells in humanized mice

We examined the reconstitution of human αβ and γδ T cells in huMice using multicolor flow cytometry (**Fig. 1**). The gating scheme is shown for a representative sample of PBMC derived from a huMouse (**Fig. 1A**). We compared these results to PBMC samples from HIV seronegative humans, with data from a representative donor is shown in **Fig. 1B**. We observed a high level of reconstitution of human CD45^+^ cells (∼90%) in the peripheral blood of huMice (**Fig. 1C**). Approximately 90% of these human CD45^+^ cells were CD3^+^ T cells, of which on average were comprised of 80% CD4^+^T cells and 15% CD8^+^ T cells (**Fig. 1D**). This CD4/CD8 ratio was slightly higher than what is typically seen in humans, as shown with the 4 donors we tested that displayed a mean of 70% CD4^+^ T cells and 30% CD8^+^ T cells (**Fig. 1D**). We also analyzed the γδ T cell subsets present in the peripheral blood of huMice and determined a mean of 0.3% and 0.7% of total CD3^+^ T cells being comprised of Vδ1 T cells and Vδ2 T cells respectively (**Fig. 1E**). The relative frequencies of these two subsets are comparable to, albeit lower than, the γδ lymphocyte populations found in the peripheral blood of healthy humans represented in our analysis showing 1% and 1.6% of total CD3^+^ T cells being Vδ1 and Vδ2 T cells respectively (**Fig.1E**). To our knowledge, this is the first report to describe the reconstitution of human γδ T cells in huMice.

**Figure 1.**
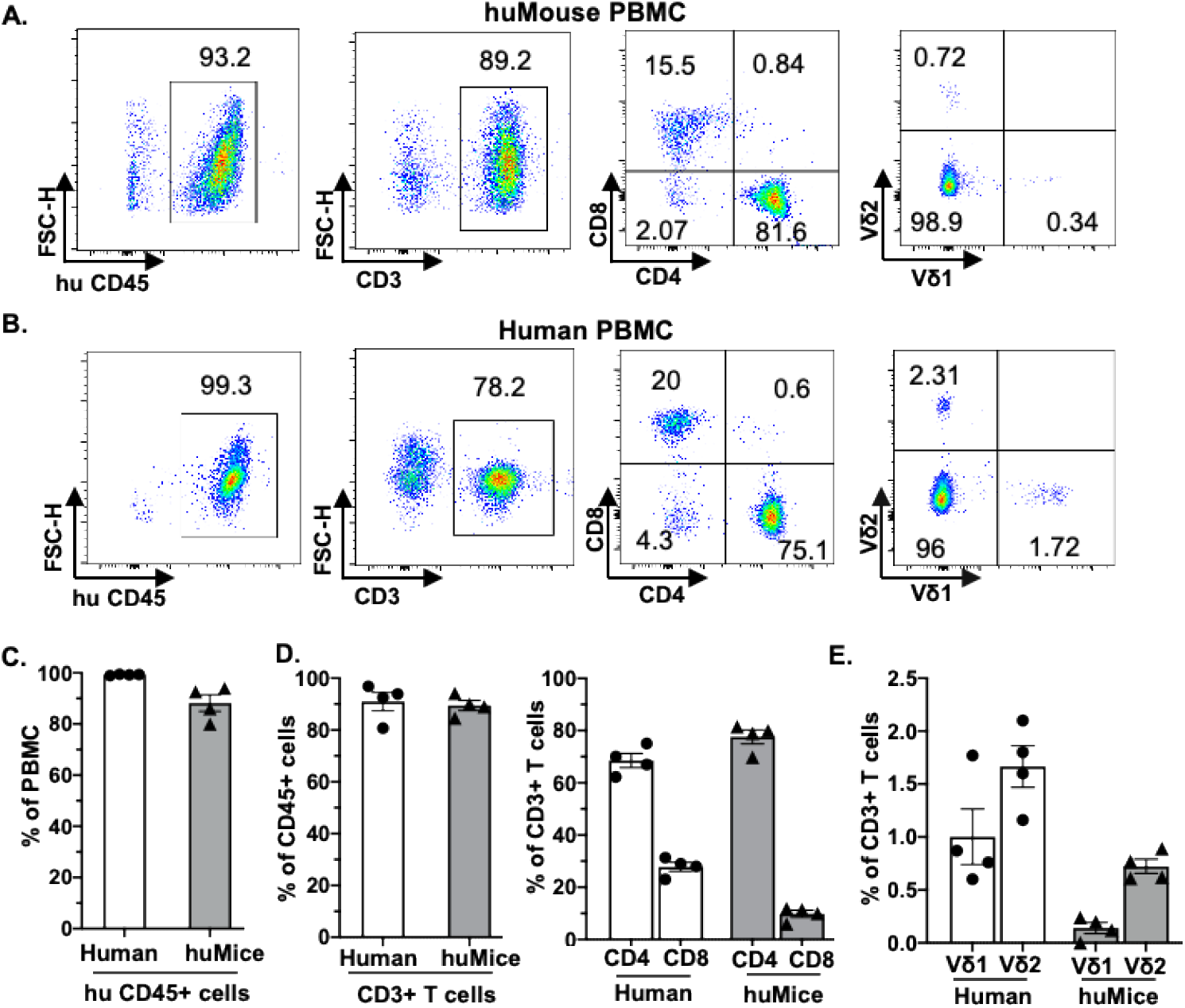
Human αβ and γδ T cell development in the peripheral blood of humanized mice. (A-B) Representative flow cytometry analysis of human immune cell (hCD45^+^) reconstitution along with lymphocytes subsets, including αβ T cells (CD3^+^), (CD4^+^), (CD8^+^), and γδ T cells (Vδ1 and Vδ2 T cell subsets) in PBMC of huMice (A) at 10 weeks after transplantation and uninfected human (B). (C-E) Quantification of human CD45^+^ lymphocytes (C), human αβ (D), and γδ (E) T cell subsets in PBMCs of huMice and healthy humans (n=4).

We also examined human immune cell populations reconstituted in the transplanted human thymus and endogenous murine spleen of huMice (**Fig. 2**). The gating scheme is shown for a representative sample of immune cells isolated from the human thymus (**Fig. 2A**) and murine spleen (**Fig. 2D**). Of the human CD3^+^ T cells isolated from the thymic tissue, an average of 22% were CD4^+^ T cells, 16% were CD8^+^ T cells, and 60% had an immature T cell phenotype being positive for both CD4 and CD8 (CD4^+^/CD8^+^, double-positive) (**Fig. 2B**). In the murine splenic tissue, on average, the total T cell population was comprised of 80% CD4^+^ T cells and 16% CD8^+^ T cells (**Fig. 2E**). Human γδ T cell subsets were also detected in these lymphoid tissues. From the human thymus an average of 1.5% of the T cells had a Vδ1 cell phenotype and 0.2% were Vδ2 T cells (**Fig. 2C**). We observed a slightly higher prevalence of γδ T cell subsets isolated from murine spleen tissue, with a mean of 2.2% and 0.9% of the total T cell fraction consisting of Vδ1 T cells and Vδ2 T cells respectively (**Fig. 2F**). We observed that Vδ2 T cells were predominantly present in the peripheral blood of huMice (**Fig. 1D**), while Vδ1 T cells were predominantly present in the lymphoid tissues of huMice (**Fig. 2C, F**). The endogenous murine spleen of the huMouse, hereafter referred to as the humanized spleen, had an approximate 2-fold higher reconstitution of γδ T cells than what was found in the thymus. This overall distribution of γδ T cell subsets in huMice is comparable to those in human peripheral blood and tissue [13, 14]. In summary, these findings demonstrated that huMice sustain physiologically similar proportions of human αβ and γδ T cells in the periphery, transplanted human thymus, and endogenous (humanized) murine spleen.

**Figure 2.**
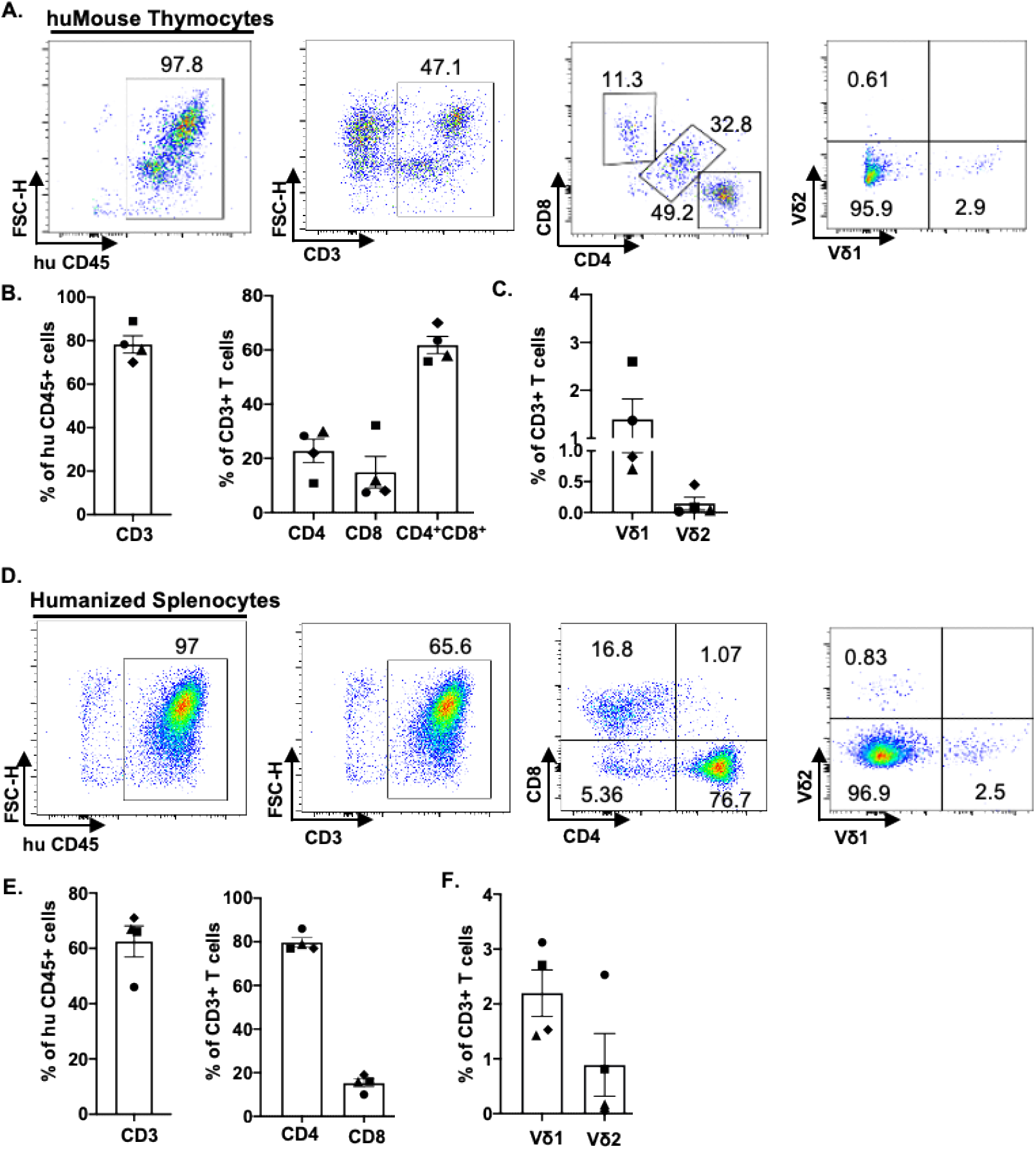
Human αβ and γδ T cell development in lymphoid tissues of a humanized mouse model. (A & D) Representative flow cytometry analysis of human immune cell (hCD45^+^) reconstitution along with lymphocytes subsets including αβ T cells (CD3^+^), (CD4^+^), (CD8^+^) and γδ T cells (Vδ1 and Vδ2 T cell subsets) in lymphoid tissue [thymus (A) and murine spleen (D)] of huMice (A) at 22 weeks post-transplantation. Quantification of human αβ and γδ T cells in the engrafted human thymus (B & C) and murine spleen tissue (E & F) of huMice at 22 weeks post-transplantation (n = 4).

### HIV infection alters γδ T cell populations in humanized mice and humans

To investigate the impact of HIV infection on γδ and αβ T cell populations, we infected huMice reconstituted with a CCR5-tropic laboratory strain of HIV-1 (NL4-3). We have previously shown HIV replication kinetics in huMice [15], which is similar to HIV replication kinetics in adult humans [16]. Consistent with the previous studies, HIV RNA copies were detected in the peripheral blood of the HIV-infected huMice as early as 2 weeks post-infection (**Fig. 3A**) [12, 15].

**Figure 3.**
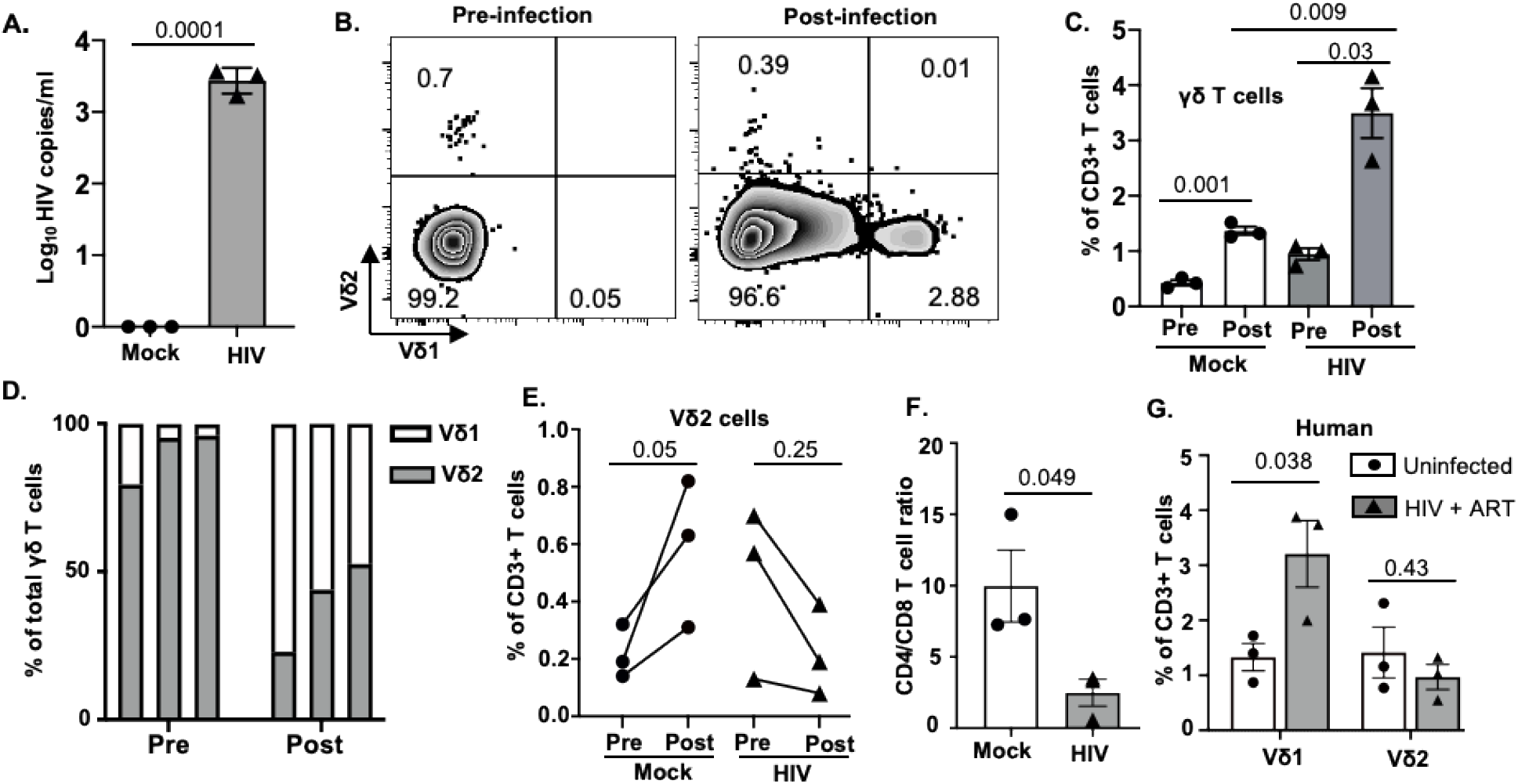
Peripheral blood γδ T cell number is altered in HIV-infected humanized mice and humans. (A) HIV-1 replication (HIV RNA genome copies per ml) in the blood following HIV_NL4_ inoculation at 1X 10^5^ IU per mouse measured by qPCR (n = 3 per group). (B) Frequency of total γδ T cells before and after HIV infection in mock and HIV-infected huMice analyzed by flow cytometry (n = 3 per group). (C) Representative flow plot showing the change in frequency of γδ T cell subsets before and after HIV infection. (D) Graphical representation of the change in frequencies of Vδ1 and Vδ2 cells within γδ population pre- and post-HIV infection. (E) Quantitation of changes in Vδ2 T cell frequency pre- and post-HIV infection in peripheral blood of HIV-infected and non-infected humanized mice. (F) Comparison of changes in CD4^+^/CD8^+^ T cell ratio in peripheral blood of HIV-infected and non-infected huMice analyzed by flow cytometry. (G) Frequency of Vδ1 and Vδ2 T cell subsets in peripheral blood of ART-treated HIV-infected and non-infected MACS participants analyzed by flow cytometry. Data are presented as a mean value ± SEM. p values <0.05 were considered statistically significant. P values were determined using paired 2-tailed Student’s t-test for comparing changes in γδ T cells population within the same cohort at two different time points, whereas an unpaired, 2-tailed Student’s t-test was used to compare differences between 2 groups.

PBMC from mock-inoculated and HIV-infected huMice were collected before and after HIV infection for further viral load analysis, and these mice were sacrificed for tissue collection 4 weeks after infection. We first determined the proportion of γδ T cells present in PBMC of HIV-infected and non-infected huMice before and after HIV infection.Representative flow cytometry analysis plots displaying the percentage of γδ T cells present at pre-and post-infection time points are shown in **Fig. 3B**. We noted an approximate 3-fold increase in total γδ T cell population in peripheral blood of both HIV-infected (p=0.03) and mock-infected huMice (p=0.001) at the post-infection time point as compared to pre-infection (**Fig. 3C**).. Interestingly we found that post-infection peripheral blood total γδ T cell frequency was approximately 2.3-fold higher in HIV-infected huMice than non-infected huMice [p=0.009] (**Fig. 3C**). This suggests that HIV infection elevates the total γδ T cell population in huMice.

Next, we determined the impact of HIV infection on the frequency of γδ T cell subsets. We found that HIV infection leads to an approximate 2-fold depletion of peripheral blood Vδ2 T cells of huMice during early infection, while the Vδ1 T cell frequency increased approximately 3-fold (**Fig. 3D**). The observed decrease of Vδ2 T cells in huMice with HIV-infection was not seen in the mock-infected mice, which actually showed an increase in Vδ2 T cell frequency (**Fig. 3E**), suggesting that HIV infection causes the selective elimination of the Vδ2 cells. Furthermore, depletion of peripheral blood CD4^+^ T cells of the HIV-infected huMice results in a dramatic decrease in the CD4^+^/CD8^+^T cell ratio (**Fig.3F**). These results are consistent with what has been previously reported in human γδ T cell studies [17, 18], and similar to the trend we find among HIV-infected MACS participants on ART who display higher frequencies of Vδ1 cells and lower percentages of Vδ2 cells (**Fig. 3G**).

In HIV-infected humans, lymphoid tissues are known to be sanctuaries for the latent HIV reservoir during ART [19]. Therefore, we assessed the impact of HIV infection on the lymphocytes derived from lymphoid tissues of huMice by flow cytometry analysis. A representative gating strategy used for this analysis is shown in **Fig. 2**. Although not statistically significant, we observed an approximately 3-fold increase in the frequency of Vδ1 T cells in the human thymus (p = 0.058), and approximately 2-fold increase in the humanized spleen of HIV-infected huMice (p = 0.065) when compared to respective tissues from non-infected huMice (**Fig. 4A, B**). This suggests that the frequency of Vδ1 T cells is increased in lymphoid tissue of huMice during HIV infection. We did not find a significant difference between the Vδ2 T cell population frequencies derived from the lymphoid tissues of HIV-infected or mock-infected huMice. Besides γδ T cells, we found approximately a 2-fold increase in the proportion of cytotoxic CD8+ T cells derived from thymus and humanized spleen tissue of HIV-infected huMice as compared to the mock-inoculated mice, suggesting a rapid proliferation of cytotoxic T cells in response to HIV infection (**Fig. 4A, B**).

**Figure 4.**
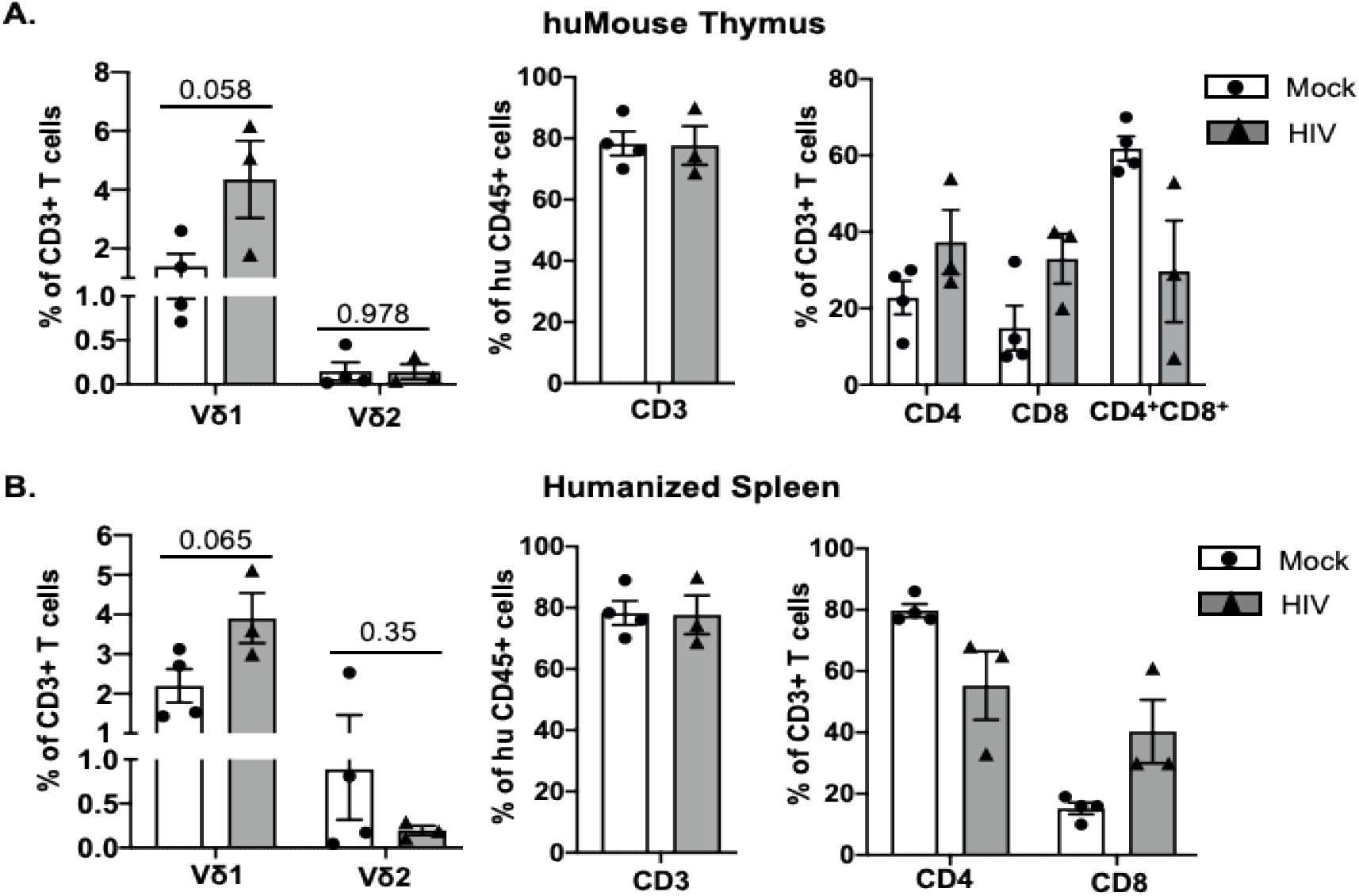
T cell number is altered in lymphoid tissue of HIV-infected humanized mice. (A-B) Quantification of human T cell subsets, γδ T cells and αβ T cells in human thymus and humanized spleen tissue of HIV-infected (n=3) and non-infected (n = 4) huMice at 22 weeks post-transplantation. Data are presented as mean values ± SEM. P values <0.05 were considered statistically significant as determined using an unpaired, 2-tailed Student’s t-test.

### Selective in vitro expansion of Vδ2 T cells from HIV-infected and non-infected humanized mice

We cultured leukocytes derived from lymphoid tissue of huMice (*n* = 6), peripheral blood of ART-suppressed HIV-infected (*n* = 4), and age-matched uninfected MACS participants (*n* = 5) and stimulated them with the combination of ZOL and rhIL-2 to selectively promote the *in vitro* expansion of Vδ2 cells. The basal percentage of Vδ2 cells within the CD3^+^ population of lymphocytes was analyzed by flow cytometry, which revealed a range of inter-individual differences among uninfected donors (1.2% - 2.2%), ART-suppressed HIV-infected individuals (0.5% - 1.2%), and huMice (0.2% - 1%). Initially, when we cultured Vδ2 T cells from the peripheral blood or the lymphoid tissues of huMice in the presence of ZOL and rhIL-2, we observed minimal expansion of Vδ2 T cells but it was not optimal. Next, we supplemented the cultures with allogenic monocytes from healthy individuals and obtained higher expansion of Vδ2 T cells. Our results show that Vδ2 T cell expansion from lymphoid tissues of non-infected mice after 10 days was approximately 4-fold higher than HIV-infected mice (p=0.013) (**Fig. 5A, B**). Similarly, we expanded Vδ2 T cells from HIV-infected and non-infected MACS participants and found that Vδ2 T cell expansion was approximately 3-fold higher in non-infected individuals than those with HIV-infection (p=0.001) (**Fig. 5C, D**). This suggests that HIV infection not only depletes the frequency of Vδ2 T cells *in vivo* but it also severely impacts the ability of these cells to expand *in vitro* in response to ZOL and rhIL-2 treatment.

**Figure 5.**
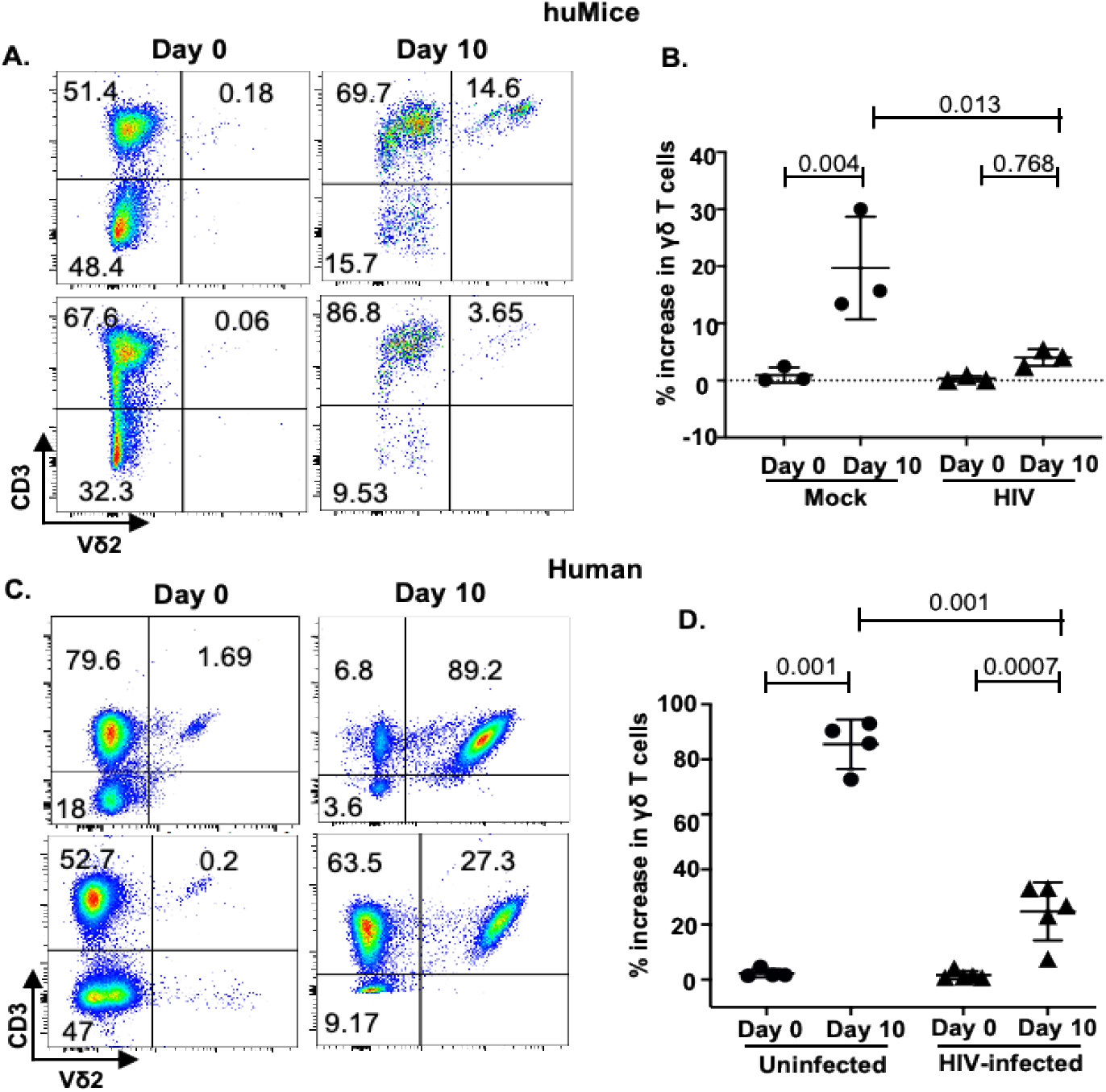
HIV infection impairs the *in vitro* expansion of Vδ2 T cells. (A-B) Splenocytes from HIV-infected/non-infected huMice were cultured in the presence of zoledronate, IL-2, and allogenic uninfected monocytes (n= 3 mice per group). (C-D) Flow plots representing *in vitro* expansion of Vδ2 cells from HIV-infected and non-infected individuals in the presence of zoledronate and IL-2. Expansion of Vδ2 cell frequency was significantly higher in non-infected donors (n=4) compared to HIV-infected donors (n=5). Data are presented as mean values ± SEM. P values <0.05 were considered statistically significant as determined using a 2-way ANOVA test.

### The phenotype of ex-vivo expanded Vδ2 T cells

The phenotype of expanded Vδ2 cells after 10 days of exposure to ZOL and rhIL-2 was analyzed in a subgroup of HIV-infected and uninfected MACS participants and HIV-infected/uninfected huMice by measuring the expression of markers of activation and differentiation by flow cytometry (**Fig. 6A**). Surface expression of the inhibitory receptor PD-1 was observed in a mean of 78% and 45% on the cultured Vδ2 cells derived from HIV-infected and uninfected huMice, respectively (p=0.04) (**Fig. 6B**). Similarly, the mean percentage of Vδ2 cells expressing PD-1 from HIV-infected and uninfected human donors was respectively 40% and 20% (p=0.001) (**Fig. 6B**). The activation markers CD69 and CD25 were co-expressed on a mean of 80% and 65% of the Vδ2 cells cultured from HIV-infected and uninfected huMice, respectively. Similarly, CD69 and CD25 co-expression was observed in a mean of 50% and 25% of the Vδ2 T cells from HIV-infected and uninfected humans, respectively (**Fig. 6C**). Together, these findings suggest that the expression of activation markers on Vδ2 cells expanded *in vitro* are slightly higher in those derived from HIV-infected humans and huMice than from their uninfected counterparts. We also evaluated the differentiation status of the cultured Vδ2 cells based on memory cell phenotypes defined as follows: (CM) central memory (CD45^−^CD27^+^), (TDM) terminally differentiated (CD45^+^CD27^−-^) and (EM) effector memory (CD45^−^CD27^−^). Although not statistically significant, we noted an increase in the TDM phenotype and a decrease in the CM and EM phenotypes in the *in vitro* expanded Vδ2 T cells derived from HIV-infected huMice compared to the Vδ2 cells cultured from uninfected huMice (**Fig. 6D**). However, in humans, we found an approximately equal distribution (20-30%) of EM, CM, TDM phenotypes between HIV-infected and non-infected individuals (**Fig. 6E**).

**Figure 6.**
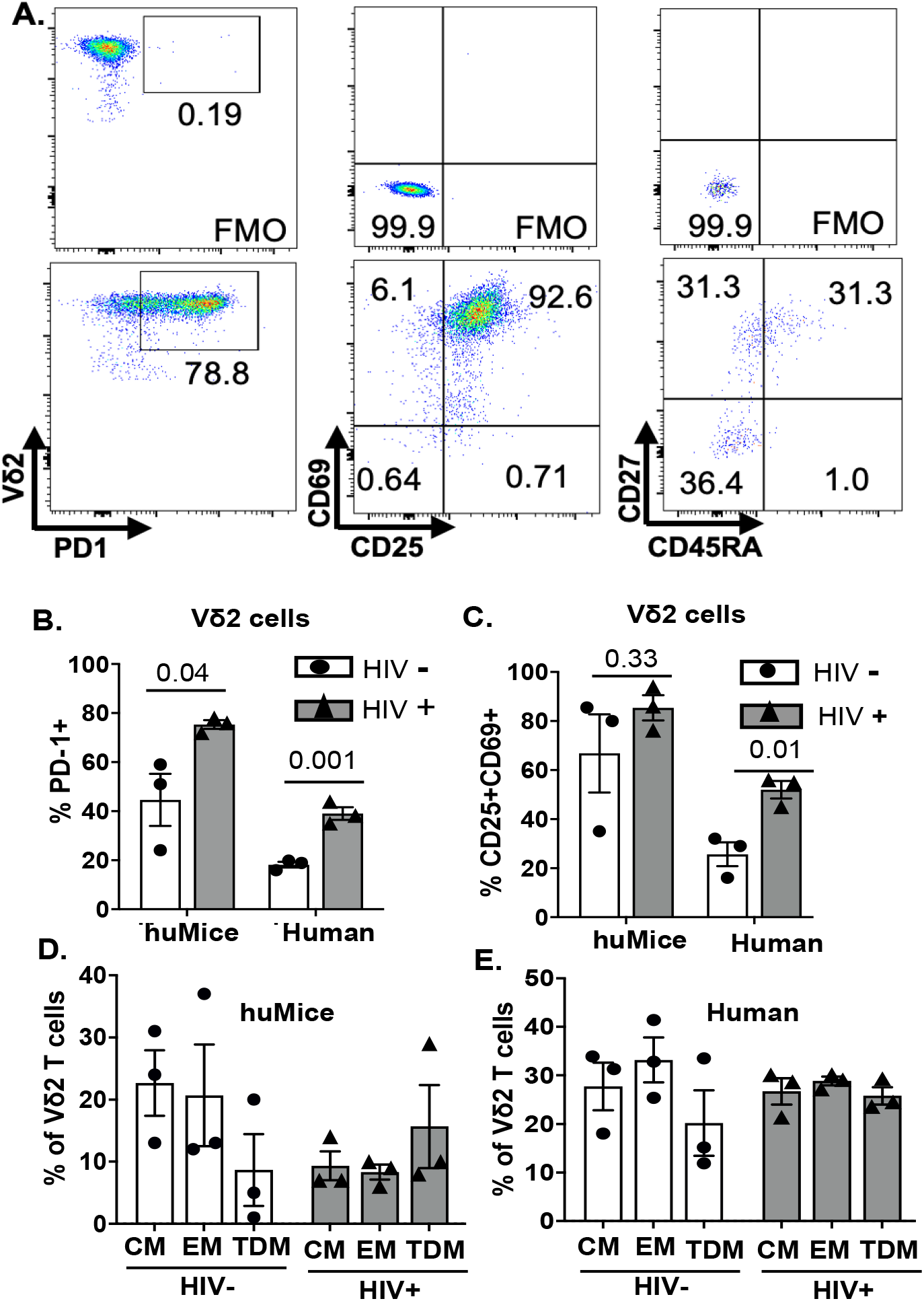
Phenotype characterization of cultured Vδ2 cells. The phenotype of Vδ2 cells from 6 huMice and 6 HIV-infected/non-infected donors after the expansion was analyzed by flow cytometry. (A) Representative flow cytometry analysis of expanded Vδ2 cells from lymphoid tissue of humanized mice expressing activation, inhibitory, and differentiation markers. (B) Expression of the checkpoint inhibitory marker PD-1 on Vδ2 cells expanded from HIV-infected and non-infected huMice and humans. (C) Dual expression of activation markers CD69 and CD25 on Vδ2 cells expanded from HIV-infected and non-infected huMice and humans. (D) Percentage of Vδ2 cells defined as central memory (CM) (CD45^−^CD27^+^), terminally differentiated (TDM) (CD45^+^ CD27^−^) and effector memory (EM) (CD45^−^CD27^−^) derived from HIV-infected and non-infected huMice. (E) Percentage of Vδ2 cells derived from HIV-infected and non-infected human MACS participants defined as having EM, CM, TDM phenotypes. Data are presented as mean values ± SEM. P values were determined using 2 tailed unpaired t-tests between the 2 groups.

### Adoptive transfer of Vδ2 T cells enhances HIV infection in humanized mice

Many *in vitro* studies have demonstrated a protective role of γδ T cells against HIV infection [8, 9, 20]. Therefore, we tested the impact of Vδ2 cells on HIV infection in the *in vivo* huMice model through the adoptive transfer of *ex vivo* expanded allogeneic Vδ2 cells. As discussed above, *in vitro* expansion of Vδ2 T cells from HIV-infected individuals was not optimal. We overcame this barrier in our adoptive transfer experiment by utilizing Vδ2 T cells expanded from allogeneic non-infected individuals. A similar strategy was previously demonstrated to be safe and effective in humans [21]. Moreover, it is therapeutically relevant because Vδ2 T cells lack functional MHC-restriction and therefore pose a minimal risk for developing graft-versus-host complications [22]. They may however serve as targets for an allogeneic response by the engrafted immune cells.

HuMice were grouped into two different cohorts: one cohort received only activated CD4^+^ T cells from HIV-infected human donor (CD4-only cohort), while the other cohort received activated CD4^+^ T cells from HIV-infected human donor as well as the cultured activated allogenic Vδ2 cells from an HIV non-infected human donor (CD4+Vδ2 cohort). Reconstitution of human Vδ2 and CD4^+^ T cells in the peripheral blood of huMice was examined via flow cytometry two weeks after the adoptive transfer procedure. We found that a mean of 50% of all T cells were Vδ2 T cells in the peripheral blood of CD4+Vδ2 cohort, whereas less than 1% of T cells were Vδ2 T cells in CD4 only cohort (p=0.03) (**Fig. 7A**), which indicated successful engraftment of human Vδ2 T cells in the huMice. Next, we confirmed HIV replication in the plasma of huMice by qPCR. Surprisingly, we observed a viral load in the CD4+Vδ2 cohort was approximately 2-fold higher than the CD4-only cohort (**Fig. 7B**) (p=0.042). Hypothesizing that this increase in viral load could be due to HIV-infection of the adoptively transferred Vδ2 T cells, we decided to analyze the CD4^+^ and Vδ2 T cell subsets in the peripheral blood of both cohorts at 2 weeks post-adoptive transfer. Representative flow cytometric plots of HIV p24 from both cohorts are shown in **Fig. 7C** and **7D**. Again, we observed an approximate 2-fold higher presence of HIV p24 in CD4^+^ T cells (p=0.020) and Vδ2 T cell (p=0.049) in the CD4+Vδ2 cohort of huMice compared to the reference CD4-only cohort (**Fig 7E, F**). Therefore, our results indicate that the adoptive transfer of Vδ2 T cells exacerbated HIV infection.

**Figure 7.**
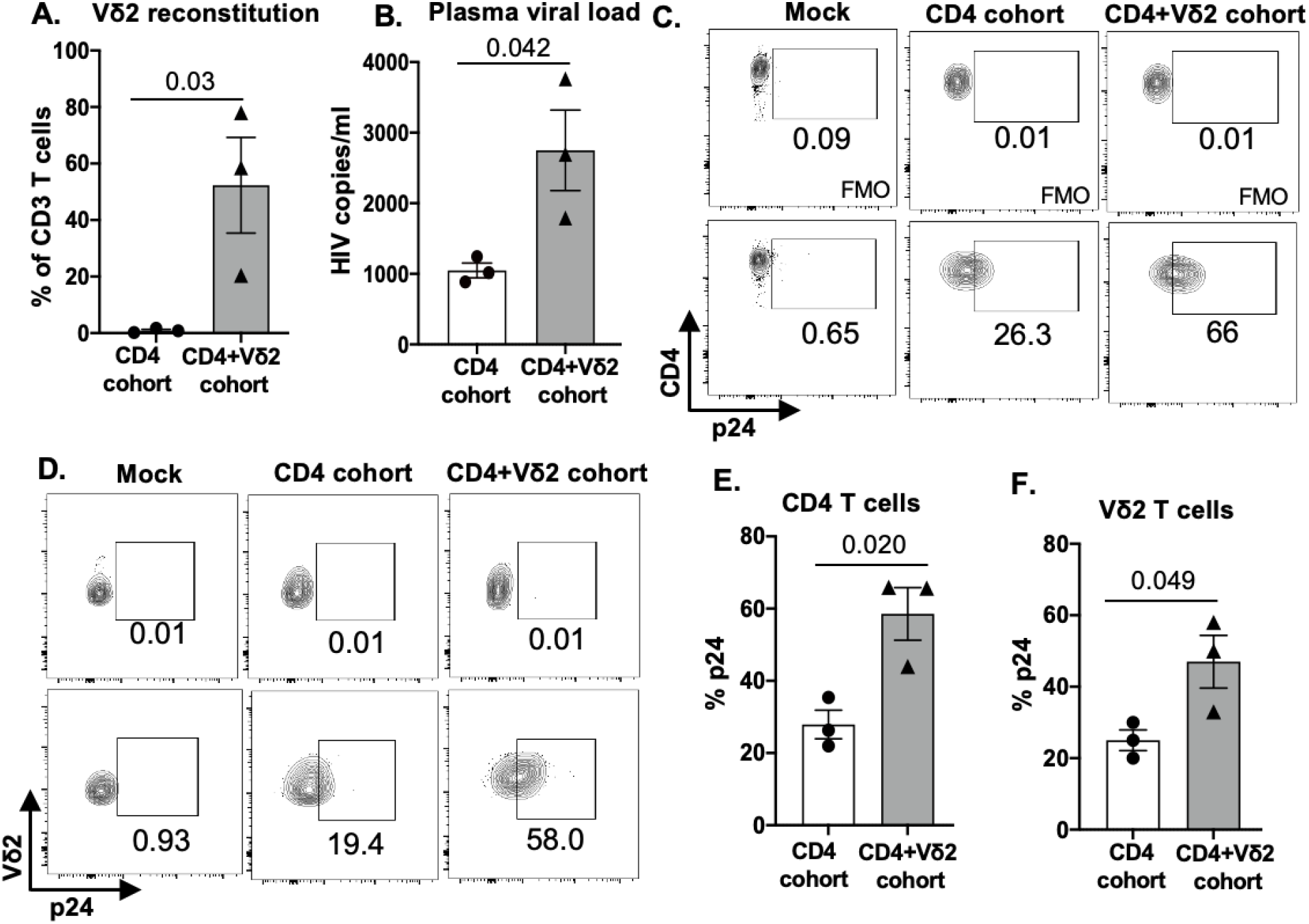
Adoptive transfer of Vδ2 T cells exacerbates HIV infection in huMice. A) Vδ2 cell number significantly increased post-adoptive transfer in peripheral blood of humanized mice (n=3 per group); analyzed by flow cytometry. (B) HIV viral load increased significantly in plasma of Vδ2+CD4-treated humanized mice as compared to CD4-treated humanized mice; measured 2 weeks post-adoptive transfer by qPCR (n=3 per group). (C & E) Representative flow cytometry analysis of peripheral blood CD4^+^ T cells and Vδ2 cells that are expressing HIV p24 respectively. (D & F) HIV p24 is significantly higher in peripheral blood CD4^+^ and Vδ2 T cells of humanized mice that received CD4+Vδ2 treatment as compared to the humanized mice that received only CD4^+^ T cells treatment respectively (n=3 per group). Data are presented as mean values ± SEM. P values were determined using 2 tailed paired t-test within the treatment groups.

Despite low or lack of CD4 receptor expression on Vδ2 T cells, our *in vivo* data suggest that these cells can indeed be targets of HIV infection. This is in accordance with a previous study from Sarabia et al, which reported that resting Vδ2 cells act as a reservoir for latent HIV infection [23]. We posited that HIV infection could impact the phenotype of Vδ2 T cells to make them more susceptible to direct infection. Since Vδ2 T cells already express high levels of the CCR5 co-receptor, we examined whether the expression of the CD4 receptor on Vδ2 T cells was induced on this cell type during HIV infection. Prior to adoptive transfer, less than 5% of endogenous and *in-vitro* cultured Vδ2 T cells expressed the CD4 receptor, but at 2 weeks after adoptive transfer, we indeed detected a mean of 30% of Vδ2 T cells expressing the CD4 receptor in both the cohorts (**Fig. 8**). Contrary to the previous reports [9, 24] highlighting the protective function of Vδ2 T cells in controlling HIV infection *in vitro*, our result suggests that HIV infection can drive CD4 expression on Vδ2 T cells *in vivo*, priming them to become targets for HIV infection and contributors to viral dissemination.

**Figure 8.**
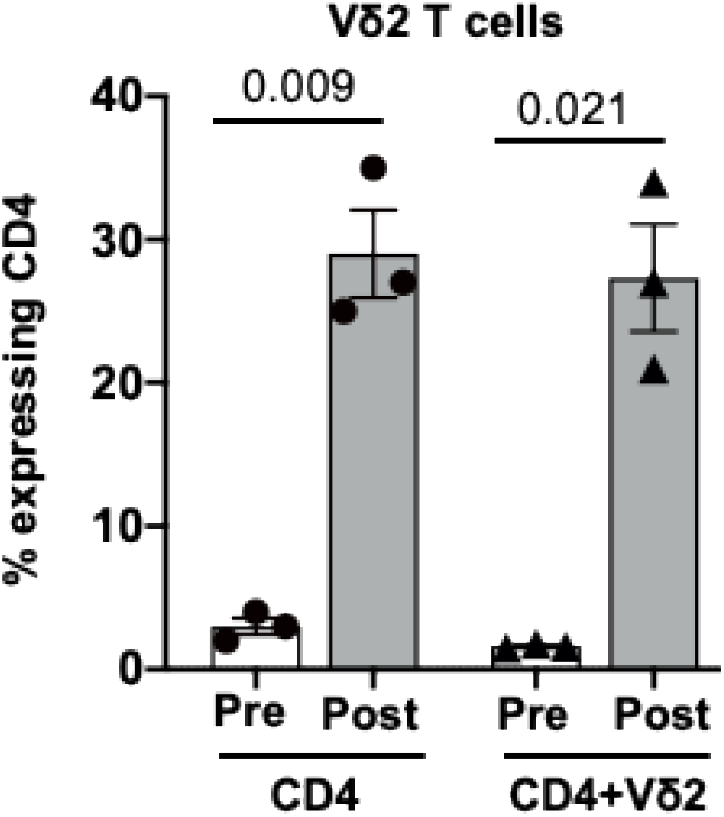
Induction of CD4 expression on Vδ2 T cells *in vivo* during HIV infection. CD4^+^ T cells from an HIV-infected MACS participant were administered to huMice with or without co-transfer of *in vitro* activated Vδ2 T cells. CD4 expression on human Vδ2 T cells from the huMice was measured by flow cytometry analysis pre-and post-(2 weeks) cell transplant. Data are presented as mean values ± SEM. P values were determined using 2 tailed paired t-test within the treatment groups.

## Discussion

γδ T cells are the first line of defense against many pathogens, but their frequency and functions are severely altered in the setting of many infectious diseases, including HIV. Despite long-term ART and viral control, γδ T cells do not reconstitute in HIV-infected individuals to their levels set prior to infection [18]. However, in HIV elite controllers, Vδ2 T cell numbers are maintained at normal levels throughout infection, implying that Vδ2 T cells play an important role in HIV infection and control. Therefore, a better understanding of their function during HIV infection will be necessary in order for them to be effectively utilized or targeted for therapeutic benefit. While prior studies have demonstrated the protective effect of γδ T cells against HIV infection *in vitro* [8, 9, 20], there is a paucity of information available and a gap in knowledge regarding their therapeutic potential *in vivo*.

In this study, we offer the first evidence that clinical trends of γδ T cell subpopulations before and after HIV infection can be modeled in BLT huMice. Immunodeficient NSG mice exhibited robust reconstitution of human immune cells, including γδ T cells, by 12 weeks post-engraftment of CD34^+^human fetal liver cells and thymic tissues. Flow cytometric analysis of human T cell subsets revealed that Vδ1/Vδ2 T cell ratios and CD4^+^/CD8^+^ T cell ratios in both the blood and lymphoid tissues of healthy BLT huMice were comparable to that seen in healthy humans. Furthermore, we observed high levels of viremia two weeks following infection, an associated depletion of Vδ2 T cells, and an expansion of Vδ1 T cells in the peripheral blood of these animals. These features suggest that BLT huMice can overcome some of the translational limitations that exist in non-human primate models of SIV, which include unremarkable changes in Vδ1/Vδ2 T cell ratios otherwise common to human HIV infection. Our study demonstrating the *in vivo* reconstitution of Vδ2 T cells in the huMouse model also provides a proof-of-concept and basis for the design of future *in vivo* studies for further evaluating the role of human γδ T cells in the setting of HIV infection as well as other chronic diseases such as cancer.

Current HIV cure strategies center on utilizing the effector functions of conventional CD8+ cytotoxic T cell lymphocytes (CTL) to kill the HIV-infected cellular reservoir following the induction of latency reversal. Unfortunately, the need to specifically stimulate or target the activation of autologous HIV-antigen specific autologous CD8^+^ T cells *ex vivo* or *in vitro* on an individual MHC/peptide-specific level, and the existence of HIV CTL escape variants within the latent reservoir has challenged the progress of this approach [25, 26]. γδ T cells offer an attractive alternative to CTL as a potential therapeutic tool to mediate anti-HIV effector functions. Their lack of MHC-restriction may provide added benefits by raising the threshold for HIV to achieve immune escape. Moreover, since they pose a reduced risk of inducing allogeneic graft rejection, they may be considered for application in allogeneic immunotherapy settings. A previous study has shown that γδ T cells mediate inhibition of HIV replication [2], but the natural scarcity of γδ T cells in tissues and circulation indicates that these cells would likely need to be expanded *ex-vivo* in order for them to have the intended therapeutic effect. Although there are numerous *in vitro* protocols for expanding γδ T cells from bulk PBMC, two major approaches can be considered for targeting γδ T cells for clinical translation. First, both ZOL and rhIL-2 can be administered to directly increase the proliferation of endogenous Vδ2 T cells [27]. The other approach would be ex-vivo activation and expansion of Vδ2 T cells for adoptive therapy. In the HIV setting, this approach is limited by the substantial loss of Vδ2 T cells that occurs during the early stages of the infection cycle, which fail to fully recover after initiation of ART. An alternative would be to harvest Vδ2 T cells from healthy donors and expand them *in vitro* using ZOL and rhIL-2 for allogeneic delivery, as has been previously reported in human cancer clinical trials [21, 28] and non-human primate models [29]. One of these cancer trials demonstrated that the adoptive transfer of haploidentical expanded Vδ2 T cells from relatives of cancer patients was safe and effective for achieving meaningful responses [21]. We attempted to culture and expand Vδ2 T cells derived from PBMC as well as lymphoid tissue of huMice using ZOL and rhIL-2. Unfortunately, while we were able to expand these huMouse derived cells *in vitro*, we were not able to collect and generate an adequate number to carry out *in vivo* studies using this method. However, when we supplemented the cultures with allogeneic monocytes from healthy individuals to enhance ZOL-induced phosphoantigen presentation, we achieved a 20-fold increase in Vδ2 T cell expansion. Importantly, this was the first reported evidence that Vδ2 T cells derived from the lymphoid tissue of huMice can indeed be expanded *in vitro*.

Our pilot study examined the therapeutic potential of adoptively transferred Vδ2 T cells in HIV infection of BLT huMice. Although previous *in vitro* studies described the protective effect of γδ T cells against HIV infection [8, 9, 20], we did not see a therapeutic benefit with the delivery of Vδ2 T cells in huMice. In fact, treatment with the activated γδ cells resulted in a significant enhancement of HIV infection as compared to the HIV-infected huMice that were not treated with the Vδ2 T cells. Our results demonstrate that during HIV infection the Vδ2 T cells can transiently upregulate surface expression of CD4 and served as viable targets for active HIV infection and dissemination. These findings raise more questions about the role of γδ T cells in the initial sequelae of HIV infection and their potential contribution to the HIV cellular reservoir as has been previously reported [23]. To our knowledge, this is the first report demonstrating that functional human γδ T cells can be robustly reconstituted in a huMice model. This small animal model provides a platform for future mechanistic studies to explore interactions between HIV and T cell subsets and, more broadly, for *in vivo* evaluation of γδ T cells and γδ T cell-based therapies in the setting of various human diseases.

## Methods and Materials

### Construction of humanized mice

Non-Obese Diabetic. Cg-Prkdcscid Il2rgtm1Wjl/SzJ (NSG) mice were obtained from the Jackson Laboratory and bred in the Division of Laboratory Animal Resources facility at the University of Pittsburgh. The mice were bred and housed under biosafety level 1, pathogen-free conditions according to the guidelines approved by the Institutional Animal Care and Use Committee and were fed irradiated chow (Prolab Isopro RHM 3000 Irradiated, catalog 5P75-RHI-W 22, PMI Nutrition International) and autoclaved water. Human fetal tissues were obtained from the Health Sciences Tissue Bank at the University of Pittsburgh and processed under biosafety level 2 conditions. Within 12 hours of receipt of fetal human liver and thymus, CD34+ hematopoietic stem cells (HSCs) were isolated from the fetal liver as previously described [30] and cryopreserved at -170° C until transplantation. Portions of the fetal liver and thymus sample were cryopreserved at -170 C until transplantation. 8 to 10-week-old NSG mice received a radiation dose of 1.50 Gray before transplantation to myoablate the animals and were immediately transferred to biosafety level 2+ animal housing. On the day of operation, the cryopreserved CD34+ HSCs and tissues were thawed, and the tissues were minced into ∼1-mm^3^ fragments, and the irradiated mice were anesthetized using 1.5-3% isoflurane. Autologous human fetal thymus and liver tissue sections were implanted under the kidney capsule, and 150,000 CD34+ HSCs were engrafted via retroorbital injection in a volume of 100 uL. Immediately following the procedure, the mice received 150uL injections of carprofen (1 mg/mL) and ceftiofur (1 mg/mL) as an analgesic and antibiotic, respectively. These injections continued once a day for 2 days for a total of three sets of injections. Successful engraftment was determined by flow cytometric analysis of human CD45 expression on blood cells of mice, now termed huMice. Mice harboring>30% of human CD45^+^ cells were used for further experiments.

### Study participants

Participants of the Pittsburgh clinical site of the Multicenter AIDS Cohort Study (MACS) were included in this study. These participants were HIV-1 infected men who were on ART for a median duration of 12.08 years, who had a median CD4^+^ T cell count of 620 cell/µl and a viral load of <50 copies/ml. Wherever mentioned blood products from age-matched HIV-negative MACS participants were used in the study. Whole blood products from HIV-1-seronegative blood donors were purchased from the Central Blood Bank of Pittsburgh. Written informed consent was obtained from participants before inclusion in the study, which was approved by The University of Pittsburgh Institutional Review Board.

### Isolation of monocytes and peripheral blood lymphocytes

Peripheral blood mononuclear cells (PBMC) obtained from a buffy coat or whole blood were isolated by standard density gradient separation using Lymphocyte Separation Medium (Corning). Monocytes were isolated from PBMC by positive magnetic bead selection (Miltenyi Biotec), and CD4+ T cells and γδ T cell subsets were isolated by negative selection (EasySep^™^ CD4 T cell, Cat #-17952 and γδ T cell isolation kit, Cat #-19255) according to the manufacturer’s specifications, and the differentially isolated cells were cultured or cryopreserved until use.

### Flow cytometry

Single-cell suspensions prepared from peripheral blood, splenocytes and thymocytes of huMice were stained with a live/dead fixable aqua dead cell stain kit (Thermo Fisher Scientific) and fluorochrome-conjugated antibodies [anti-human CD45, anti-human CD4, anti-human Vδ2, (BioLegend); anti-human CD8, CD3, PD1, HLA-DR, CD25, CD69, CD45RA, and CD27 (Becton Dickenson); and anti-human Vδ1, (Thermo Fisher Scientific)] and HIV-p24 (KC57, Beckman Coulter). Cells were fixed using 2% paraformaldehyde, and data were acquired using an LSR Fortessa flow cytometer (BD Biosciences) and analyzed using FlowJo software. Gating was done based on Fluorescence minus one (FMO).

### *In vitro* expansion of γδ T cells

huMice were sacrificed and fully developed lymphoid tissues were collected and single cells were isolated following mechanical dissociation. Cells isolated from the lymphoid tissue were cocultured with allogeneic monocytes (4:1 ratio) from HIV-seronegative human blood bank donors in the presence of nitrogen-containing bisphosphonate zoledronate (ZOL, 5uM) (Zoledronic Acid, Selleckchem, S1314) and recombinant human (rh)IL-2 (Proleukin®, 100 IU/mL; Prometheus Laboratories) for 10 days as previously described [31]. rhIL-2 (100 IU/ml) was subsequently added every 3 days. The 10 day-cultured γδ T cells were characterized by flow cytometry analysis.

### HIV infection of humanized mice

The CCR5-tropic strain of HIV-1 (NL4-3) [32] was generated by transfection of 293T cells (ATCC; ATCC CRL-3216) with a plasmid containing a full-length HIV genome and collecting the HIV containing culture supernatant. The viral titer was determined using GHOST cells (NIH AIDS Reagent Program; catalog 3942) as previously described [33]. Supernatant from uninfected 293T cells was used as a mock control. huMice were anesthetized and inoculated with mock control supernatant or HIV-1 (∼1 × 10^5^ infectious units) by i.v. injection via retroorbital delivery.

### HIV-1 genomic RNA detection

Total RNA was purified from plasma using RNA-Bee (AMSBIO). The RNA was then reverse-transcribed using TaqMan Reverse Transcription Reagents (Invitrogen) and quantitatively detected by real-time PCR using the TaqMan Universal PCR Master Mix (Invitrogen) with primers (forward primer, 5′ - CCCATGTTTTCAGCATTATCAGAA - 3′, and reverse primer, 5′ - CCACTGTGTTTAGCATGGTGTTTAA - 3′) and detection probe targeting HIV Gag gene (5′ - AGCCACCCCACAAGA - 3′) [34]. The assay sensitivity/cutoff was 10 copies/ml.

### Adoptive transfer of T cells to humanized mice

PBMC derived CD4^+^ T cells were isolated from HIV-infected MACS participants using EasySep^™^Human CD4^+^ T Cell Isolation Kit and activated overnight with Human T-Activator CD3/CD28 Dynabeads® (Life Technologies). The next day Dynabeads were separated from the CD4^+^ T cells by manual dissociation followed by magnet isolation. The activated CD4^+^ T cells were, washed, resuspended in PBS, and adoptively transferred into huMice via intraperitoneal injection (5 million cells/100µl/mouse). PBMC from the allogenic HIV non-infected donor were cultured in the presence of ZOL and rhIL-2 for 10 days to expand the Vδ2 cells. Activated and expanded Vδ2 cells were adoptively transferred to huMice via intraperitoneal injection (10 million/100µl/mouse). The huMice were divided into two treatment cohorts; one that received only activated CD4^+^ T cells from HIV-infected donor, and the other that received the activated HIV-infected CD4^+^ T cells as well as *in vitro* expanded allogenic Vδ2 cells.

### Statistics

Differences between HIV-infected/uninfected humans and huMice were compared using the two-tailed unpaired Student t-test. Differences among the human or huMice groups were compared using the two-tailed paired students t-test. The normality of the samples was tested using the Shapiro-Wilk normality test. Statistical analyses were performed using the Prism8 (GraphPad Software) and p values <0.05 were considered statistically significant. The sample numbers and statistical analyses used are specified in each figure legend.

### Approval for acquisition and use of human fetal tissue and biological agents

We described the approval of use of human fetal tissue and biological agents in the previous study [15]. Briefly human fetal liver and thymus (gestational age of 18–20 weeks) were obtained from medically or elective indicated termination of pregnancy through Magee-Women’s Hospital of UPMC via the University of Pittsburgh, Health Sciences Tissue Bank and Advanced Biosciences Resources, Inc.. Written informed consent of the maternal donors was obtained in all cases, under IRB and federal/state regulations. The use of human fetal organs/cells to construct huMice was reviewed by the University of Pittsburgh’s IRB office, which has determined that this submission does not constitute human subject research as defined under federal regulations (45 CFR 46.102 [d or f] and 21 CFR 56.102[c], [e], and [l]). The use of human hematopoietic stem cells was reviewed and approved by the Human Stem Cell Research Oversight (hSCRO) at the University of Pittsburgh.

### Approval for use of animals and biological agents for *in vivo* experiments

The use of biological agents (e.g., HIV), recombinant DNA, and transgenic animals was reviewed and approved by the Institutional Biosafety Committee (IBC) at the University of Pittsburgh. All animal studies were approved by the IACUC at the University of Pittsburgh and were conducted following the NIH guidelines for housing and care of laboratory animals as well as the ARRIVE guidelines 2.0 for reporting of *in vivo* experiments involving animal research [35].

## Author Contributions

S.B., M.T.B, and R.B.M. contributed to the experimental and study design. S.B. and Y.A. performed the experiments. S.B. analyzed the data and prepared the manuscript. Y.A., R.B.M., C.R.R., M.T.B, and M.T.L. contributed to the interpretation of the results and critically edited the manuscript. All authors have read and agreed to the published version of the manuscript.

